# Ablation of SAMD1 in Mice Causes Failure of Angiogenesis, Embryonic Lethality

**DOI:** 10.1101/2022.01.11.473462

**Authors:** Bruce Campbell, Sandra Engle, Terence Ozolins, Patricia Bourassa, Robert Aiello

## Abstract

Pathological retention of LDL in the intima is involved in atherosclerosis, although the retention mechanisms are not well-understood. Previously, we reported Sterile Alpha Motif Domain Containing 1 (SAMD1), a protein secreted by intimal smooth muscle cells in atherosclerotic lesions, appears to bind LDL in extracellular matrix around intimal cells. Fab-fragment inhibitors of apparently irreversible SAMD1/LDL binding reduced LDL retention in carotid injury models, but did not have a significant effect on early spontaneous lesion development. Our histology of mouse atherosclerosis models revealed extensive SAMD1 expression, possibly related to cell phenotype modulation and antigen presentation. Although the normal function of SAMD1 is unknown, it may have multiple epigenetic roles. For this report, we generated SAMD1−/−, SAMD1−/+, and SAMD1−/+ apoE−/− mice to further explore SAMD1’s role in atherosclerosis. SAMD1 was found in tissues throughout the SAMD1+/+ and SAMD1−/+ embryos. Homozygous loss of SAMD1 was embryonic lethal: at embryonic day 14, organs were partially developed and/or degraded; heads and brains were malformed; no blood vessels were observed; red blood cells were scattered and pooled, primarily near the embryo surface; and cell death was occurring. Development appeared normal in heterozygous SAMD1 embryos, but postnatal genotyping showed a reduced ability to thrive. Growth of atherosclerotic lesions in SAMD1−/+ apoE−/− after 35 weeks was not significantly less than in mice SAMD1+/+ apoE−/− mice.

## Introduction

Human SAMD1, accession number Q6SPF0, is a 538 a.a. protein mapping to chromosome 19p13.12, and mouse SAMD1, accession number D3YXK1, is a 519 a.a. protein on chromosome 8, 40.24 cM. SAMD1 has homology across primates, canines, rodents, and rabbits, and orthology with at least 250 organisms, and SAMD1 RNA is found in numerous organs and tissues in humans and mice [1], [2], [3]; the extensiveness of expression suggests roles in adult homeostasis. SAMD1 was minimally studied until very recently [4]. A 2021 report supports SAMD1’s participation in LDL retention and uptake, finding that SAMD1 knockdown suppresses Vascular Smooth Muscle Cell (VSMC) differentiation and proliferation, and that SAMD1/LDL binding on VSMCs leads to LDL oxidation and foam cell development [5]. Another 2021 report concludes that SAMD1 interacts with CpG islands and chromatin regulators to mediate transcriptional repression and is required for proper embryonic stem (ES) cell differentiation [6]. Earlier reports suggest broad epigenetic functions for SAMD1:

- In a study of regulatory regions that can determine cell identity, SAMD1 was the strongest enhancer in H3K4me3 [7];
- SAMD1 was among the highest 5% of proteins for association with both TAF3 recognized H3K4me3-modified, and bivalently H3K4me3/H3K27me3-modified chromatin. TAF3 is a regulator of p53 which modulates cell cycle arrest and programmed cell death [8];
- Oscillatory cAMP signaling rapidly alters H3K4 methylation; SAMD1 is listed as a gene in which promoter peaks were rapidly down-regulated by cAMP and then returned to baseline quickly after cAMP washout [9].
- It can be inferred that SAMD1 is a methyl-lysine reader which recognizes methylated lysine residues on histone protein tails, and are associated with repression of gene expression [10];
- SAMD1 may be involved in myogenic and adipogenic cell fate commitment [11], and changing cell transcription response to external stimuli [9].
- SAMD1’s participation in epigenetic regulation may be related to changes in cell phenotypes, and is reported to be a participant in H3K4-methylation factors. H3K4me3 is a permanent epigenetic marker of SMC lineage in the SMMHC promoter [12].

A recent consensus statement concludes that LDL is causal in atherosclerosis [13]. In normal intima, LDL continuously and rapidly perfuses through the artery wall, entering through the endothelial layer and exiting through the lymph system [14], but retention of LDL in the intima is widely believed to initiate and continue lesion development [15]. SAMD1 was originally isolated from rabbit injury-model atherosclerotic lesions, which are rich in SMCs, by screening for low density lipoprotein (LDL) binding [16]. In mouse models of atherosclerosis, the initial retention mechanism may be proteoglycan/LDL binding, which triggers inflammatory responses leading to monocyte recruitment into the intima and conversion of monocytes to macrophages. Medial SMCs dedifferentiate, migrate into the intima, where they proliferate, differentiate into a macrophage-like phenotype, and retain LDL in their extracellular matrix (ECM). Like macrophages, these SMCs ingest and lyse LDL, and become foam cells. In mouse models, cells of SMC origin, but lacking typical SMC morphology and markers, may represent the majority of foam cells by 20-30 weeks [17].

SAMD1/apoB distribution patterns in apoE−/− and LDLR−/− mice observed with IHC at 20 and 34 weeks showed localization in endothelial cells (ECs), and at the cell surface of, and within, foam cells presumably of SMC origin. Confocal microscopy at 45 weeks showed SAMD1/apoB colocalization at EC cell surfaces, in SMC ECM, and in foam cells in LDLR−/− mice [3]. Further, SAMD1 distribution patterns in mouse arteries appeared to correlate with EC, SMC, and macrophage phenotype changes; for example, less SAMD1 was seen in presumably dedifferentiating medial SMCs and more was seen in presumably differentiating intimal SMCs [3]. In human lesions, confocal microscopy established both SAMD1 in ECs, and SAMD1/apoB colocalization in the thick ECM surrounding SMCs and encircling lipid droplets that fill foam cells [16].

We hypothesized that SAMD1/LDL binding may be involved in lesion growth, but only when SMCs are present in the intima [3], [16]. LDL retention in mouse carotid injury model lesions, which are also rich in SMCs [18], was greatly reduced by *in vivo* injection of systemic SAMD1/apoB binding inhibitors, but reduction in spontaneous lesion initiation in apoE−/− mice after 12 weeks on modified western diet, was non-significant, possibly due to insufficient SMC content [3]. Thus, SAMD1 translocated to the ECM by a macrophage-like SMC phenotype may be responsible for binding LDL in the intima after lesion initiation [3], [9]. The atherosclerotic role of SAMD1 in ECs remains unclear.

Here we report on the deletion of the SAMD1 gene in mice. Fewer homozygous embryos were observed at ED14, and none after ED18; development appeared grossly normal at ED12, but embryos exhibited numerous defects at ED14, including lack of blood vessels. Many heterozygotes, exhibiting some hormonal changes, survived past week 3 postnatal. Atherosclerotic lesion size at 35 weeks in SAMD1+/− apoE−/− mice was not affected by reduced SAMD1 mRNA expression.

## Results

### SAMD1−/− Phenotype Compared to Wild Type

The SAMD1 gene was deleted from c57bl/6 mice by recombineering (Supplementary Fig. 1). Ratios between the wild type (WT, aka SAMD1+/+), heterozygote (HET, aka SAMD1+/−), and knockout (KO, aka SAMD1−/−) mice were close to the expected Mendelian ratios through ED12, but from ED14 to ED18, there was trend towards fewer KO embryos (p<0.096) (Table 1). Dying pups were not seen, so the few KO mice that had not been resorbed by ED18 died just prior to birth. SAMD1 mRNA was not found in Mouse Embryonic Fibroblasts (MEFs) raised from KO mice; vascular cell adhesion molecule-1 (VCAM1) and Protein Kinase CAMP-Activated Catalytic Subunit Alpha (PRKACA) were substantially overexpressed in the KO compared to the WT MEFs (Fig. 1). Expression of X-Box Binding Protein 1 (XBP1), which is widely expressed during embryonic development [19], and its isoform XBP1s were non-significantly higher in the HET and KO than in the WT MEFs. SAMD1 mRNA was widely expressed in the WT throughout the conceptus; measurements taken ED8 – ED18 showed up- and down-regulations, with the highest levels expressed in the head at ED12, and the spleen at ED18 (Fig. 2).

**Figure 1:**
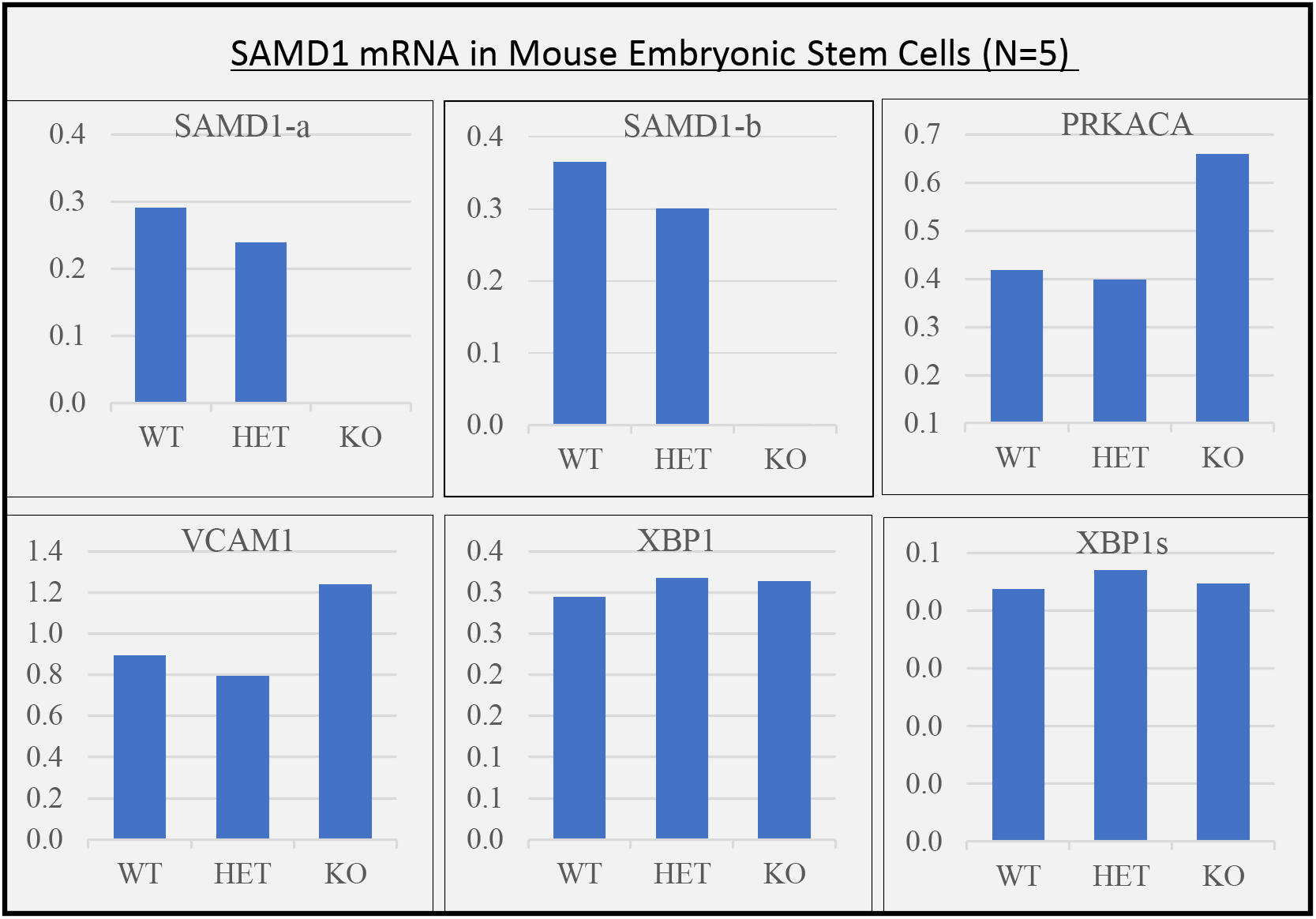
KO MEFs express no SAMD1 mRNA; HETs express less SAMD1 than WTs; KOs express more PRKCA and VCAM1 than WTs or HETs. RT-qPCR was used to quantify relative mRNA expression by Mouse Embryonic Stem Cells (MEFs) of the following proteins: SAMD1; Protein Kinase CAMP-Activated Catalytic Subunit Alpha (PRKACA); Vascular Cell Adhesion Molecule-1 (VCAM1); X-Box Binding Proteins 1 and 1s (XBP1 and XBP1s). SAMD1-a and SAMD1-b represent different primers. Note that the scales are not the same size in all 6 charts.

**Figure 2:**
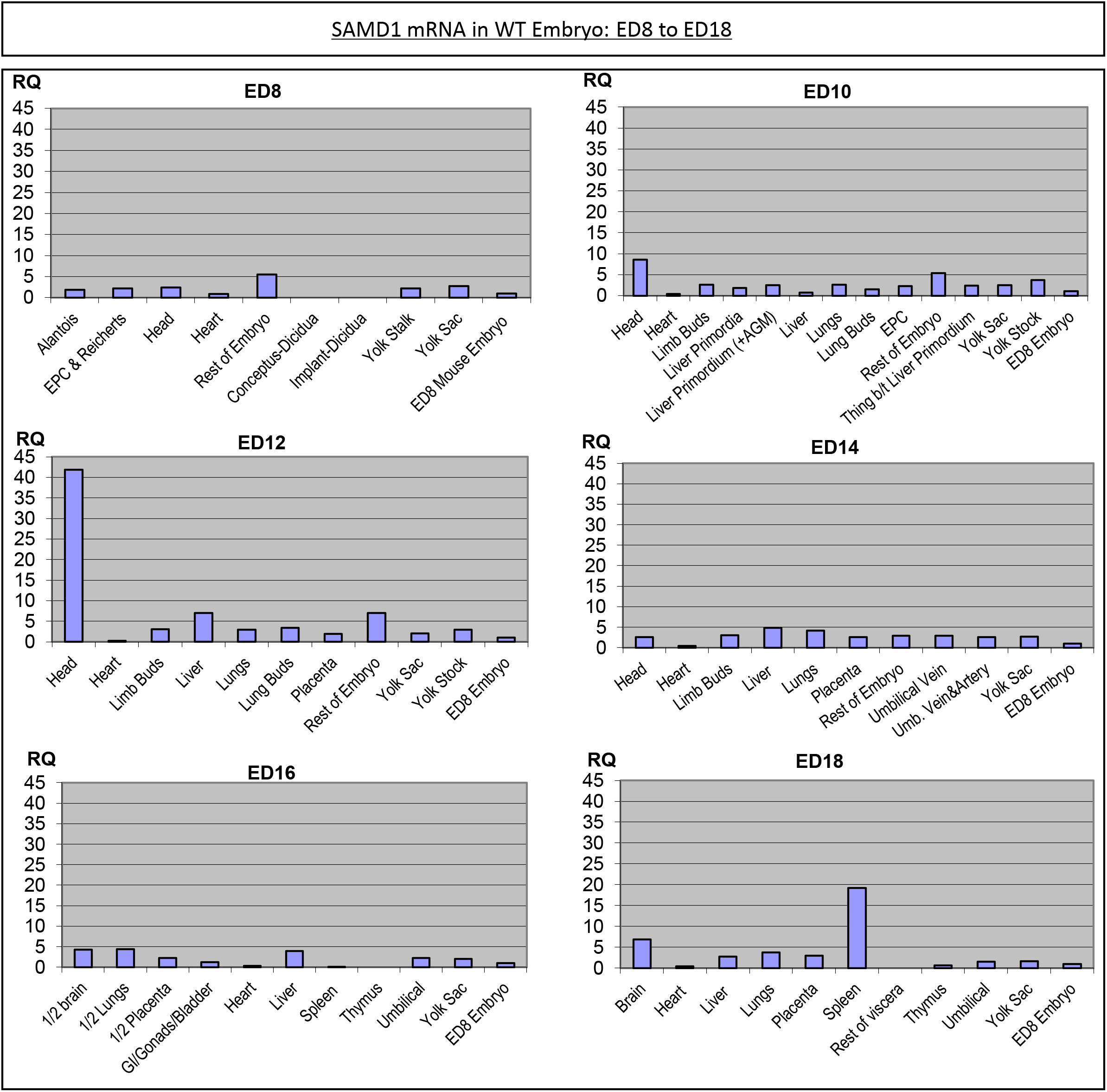
SAMD1 is expressed throughout the WT during embryonic development. RT-qPCR was used to quantify relative SAMD1 mRNA expression levels in various WT embryo tissues every other day from ED8 to ED18. As seen in the right-hand corner of each graph, levels are normalized to the ED8 embryo to better show when and where changes are occurring.

**Table 1:** Embryonic lethality may begin between ED12 and ED14. A) Embryos were genotyped between ED10 and ED18; the decreased total number of KO embryos from ED14 to ED18 suggests embryonic lethality (p<0.095). B) Additional mice were genotyped to compare ED12 to ED14. The ratios are suggestive, but the chi-square (R=0.22) is insufficient to add support to a trend.

| Embryo Genotypes and Lethality Mapping |  |  |  |  |
| --- | --- | --- | --- | --- |
| Embryonic Day | +/+ | +/- | -/- | Total N |
| 10 | 1 (5%) | 10 (56%) | 7 (39%) | 18 |
| 11 | 2 (13%) | 7 (47%) | 6 (40%) | 15 |
| 12 | 18 (23%) | 44 (56%) | 17 (22%) | 79 |
| 14 | 21 (30%) | 38 (54%) | 12 (17%) | 71 |
| 16 | 2 (11%) | 10 (56%) | 6 (33%) | 18 |
| 18 | 16 (46%) | 15 (43%) | 4 (11%) | 35 |

**Table 1: Embryonic lethality may begin between ED12 and ED14.**
| Lethality Mapping |  |  |  |  |
| --- | --- | --- | --- | --- |
| Embryonic Day | +/+ | +/- | -/- | Total N |
| 12 | 9 (26%) | 17 (50%) | 8 (24%) | 34 |
| 14 | 8 (30%) | 15 (56%) | 4 (15%) | 27 |

**Table 2:**
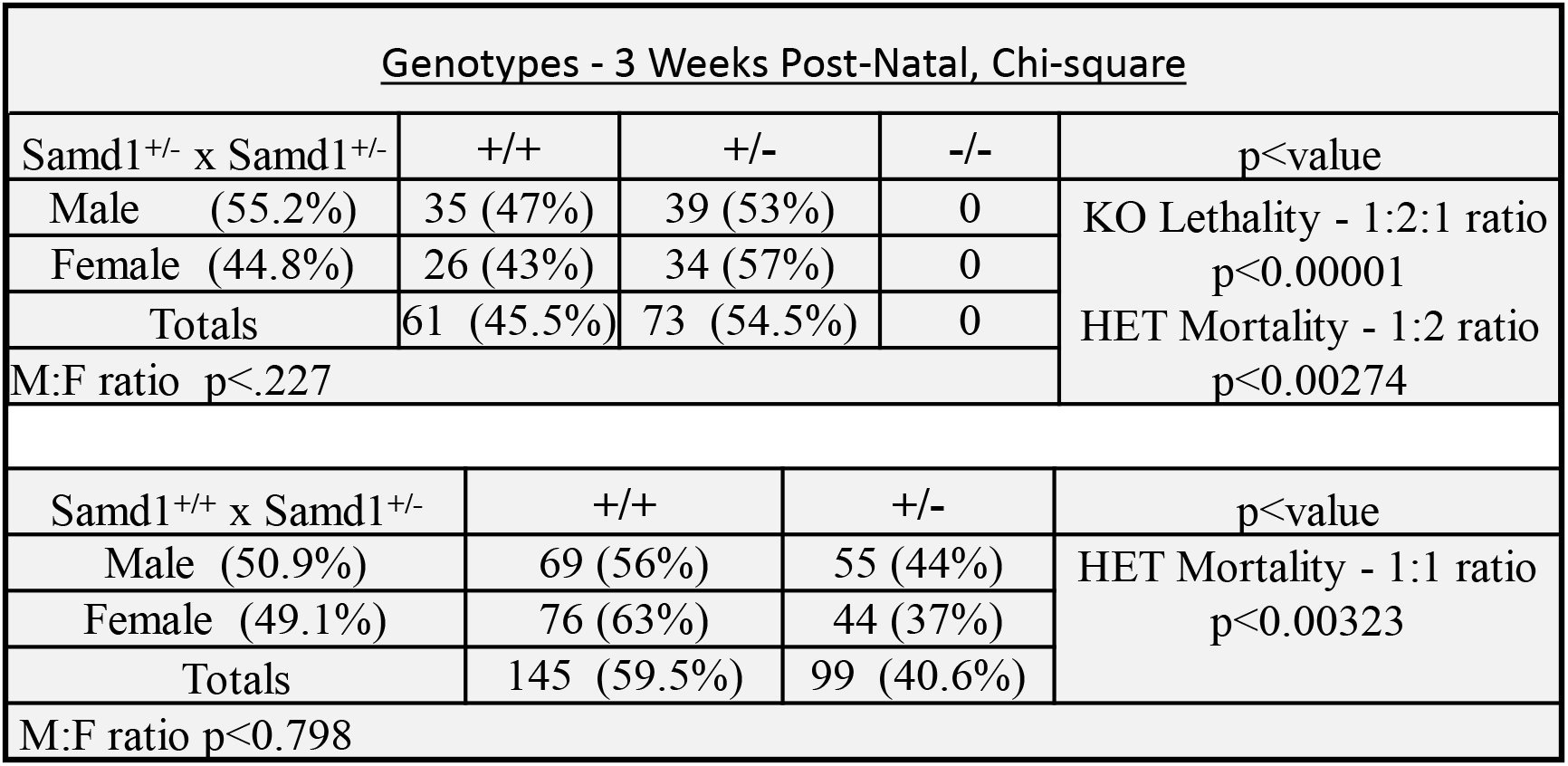
HETs may exhibit failure to thrive. No KO mice were observed three weeks postnatal, which has an essentially zero chance of occurring randomly (1:2:1 ratio). Testing for postnatal mortality, mendelian ratios for WT:HET show a high likelihood that some HETs do not survive for three weeks after parturition; results from Samd1+/− **X** Samd1+/− mice (p<0.00274; 1:2 ratio) are consistent with results from Samd1+/+ **X** Samd1+/− mice (p<0.00323; 1:1 ratio). The apparent trend towards increased mortality in HET males from the Samd1^+/−^ **X** Samd1^+/−^ group and HET females from the Samd1^+/x^ **X** Samd1^+/−^ is not significant (p<0.268).

WT and KO embryos looked grossly similar at ED12, but a broad phenotype was visible in the ED14 KOs (Fig. 3A). They were small, about the size of a WT at ED12, with shortened nasal regions, almost avascular yolk sacs, and the heads, tails, and limbs of the embryos were white. Blood vessels in the placenta were on the maternal side only, there was no vasculature on the fetal side, and blood was pooled in the yolk sac, as if blood from maternal blood vessels spilled out inside the yolk sac. Internal to the amnion, probable dorsal edema could be seen, and blood was pooled around the midsection, apparently mixed with edemic fluid. One ED14 embryo may have had a heartbeat at removal. An embryo at about ED 15.5 (Fig. 3B) lacked a skull vault, had scattered blood pooling in the head, and severe dorsal edema revealing scattered lines and spots of blood in almost transparent skin. Interestingly, the edemic fluid was clear at ED15.5, and endochondral bone development appeared to have proceeded normally in forelimbs, hindlimbs, ribs, and tail, but not the skull.

**Figure 3:**
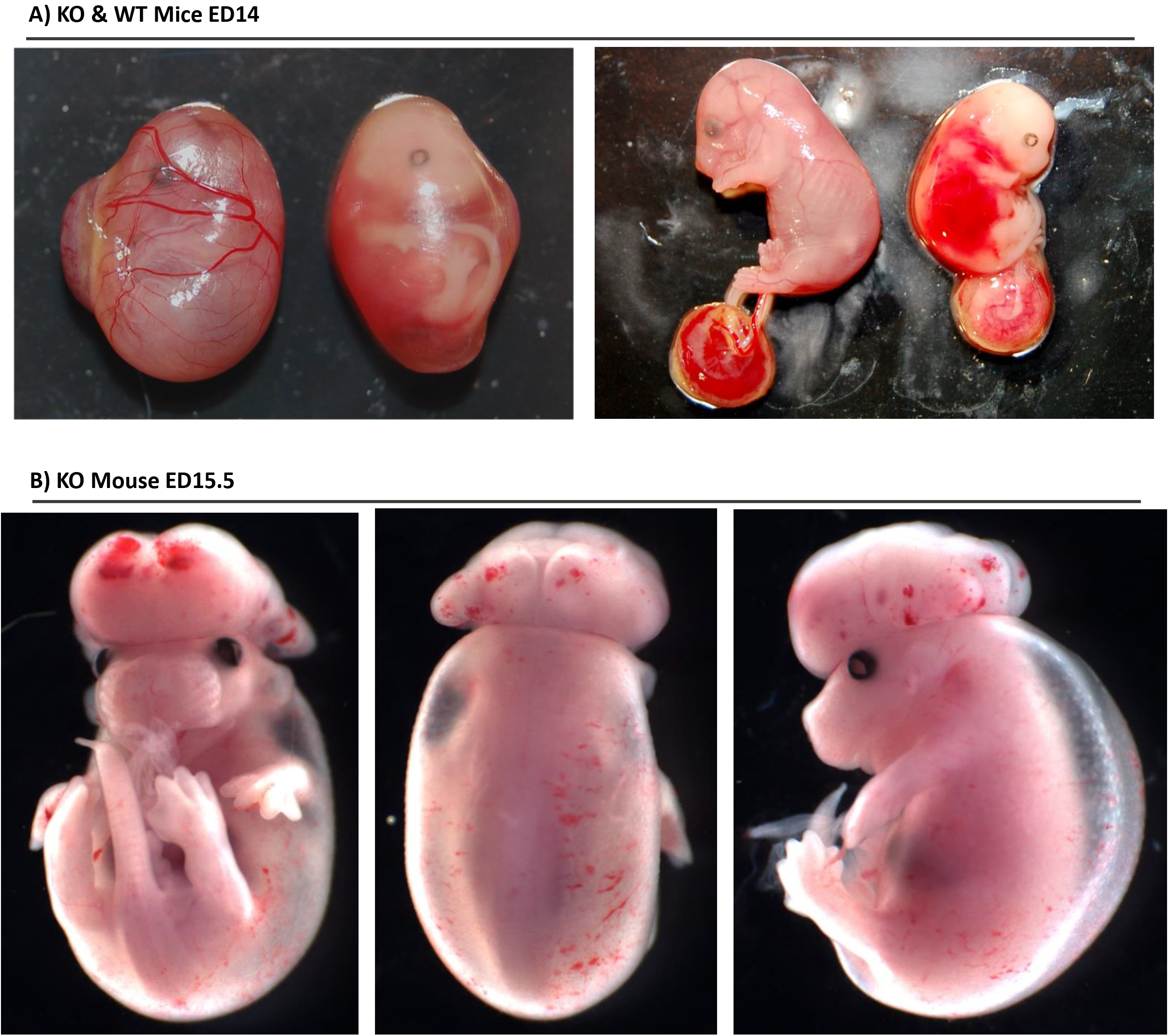
Substantial abnormalities in the KO at ED14 are visible to the naked eye. A) At ED14, WT embryo shows normal development, with branching blood vessels in the yolk sac and skin. The KO embryo yolk sac lacks blood vessels and appears to be fluid filled. With the yolk sac removed, the KO appears to be about the size of an ED12 embryo, has craniofacial abnormalities, appears to have edemic fluid mixed with blood beneath translucent skin, and has no visible blood vessels. Fore- and hind paws look grossly normal. B) At approx. ED15.5, the KO embryo has obvious exencephaly and hypertrophic brain, Severe edema is contained by transparent skin; some areas of skin have lines of red blood cells that may have once been contained in vessels, particularly above the brain. Paw and tail development appears normal.

H&E staining of the WT at ED 14 (Fig 4A) compared to the KO at ED14 (Fig 4B,C) showed abnormalities in KO embryonic organs and tissues. Internal organs were morphologically identifiable (lung), fragmented (liver), and slightly disorganized (heart). Ribs and skeletal muscle were slightly malformed; dermis was missing, possibly due to the edema causing skin loss during paraffin fixation (Fig 4C). The heart’s chambers contained immune cells but not red blood cells (RBCs), and the epithelial and pericardial layers appeared to be degrading. RBCs were not seen in the liver, or lung, but were present between two ribs; here off-center nuclei suggest immature RBCs. Eosin staining of skeletal muscle was attenuated. RBCs, just beneath where skin should be, were aligned in patterns suggesting they may have once been contained in blood vessels. There did not appear to be blood islands.

**Figure 4:**
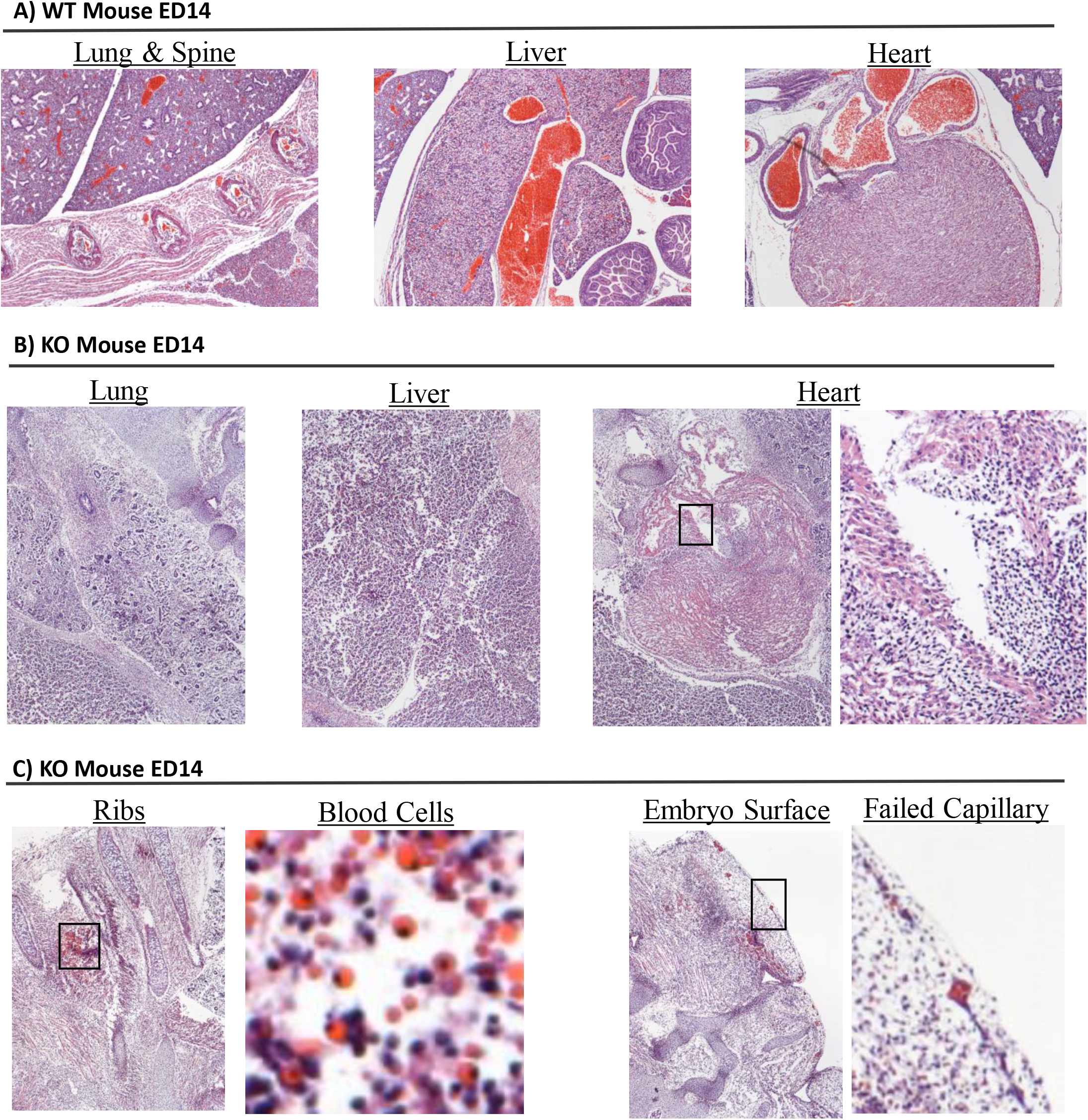
H&E staining reveals disordered organs and blood cells not contained by blood vessels. Sagittal slices of a WT and a KO mouse were H&E stained and compared. A) Samples from an ED14 WT for reference. B) Comparable samples from a KO. Lung displayed undersized and possibly defective bronchioles and alveoli. Liver was morphologically identifiable, but lacked typical structures, and separations may have occurred along failed blood vessels. Heart muscle and chambers had developed, but were fragmented, with apoptotic and early necrotic cells in the walls and muscle, and immune cells in the chambers. Blood vessels and red blood cells were absent. C) Hypertrophic chondrocytes are visible in ribs. RBCs near ribs, varying in size, some with eccentric nuclei, are not contained in blood vessels. A short distance from the embryo’s surface, aligned RBCs at the embryo surface suggest failed blood vessels. Epidermis apparently did not survive fixation in the KO.

Cluster of Differentiation 31 (CD31, aka PECAM1) and the Receptor for Vascular Endothelial Growth Factor 2 (VEGFR2, aka FLK1) are both widely used as markers for endothelial cells (ECs). To determine whether angiogenesis had occurred, samples from ED14 WT and KO mice were IHC stained with antibodies to CD31 or VEGFR2. In the WT, both probes stained capillaries and larger blood vessels throughout the mouse in the expected linear and circular patterns (Fig. 5). In contrast, in the KO heart, CD31 and VEGFR2 stained EC nuclei, cytoplasm, and fragments that together formed small broken circles and lines suggesting incomplete and/or failed blood vessels. CD31 and VEGFR2 stained epithelial cells lining the WT heart’s chambers; while in the KO, staining appeared to show endocardial fragmentation. CD31 and VEGFR2 stained apparent capillary ECs leaking cytoplasm in the KO lung and liver, and VEGFR2 faintly stained possible macrophage engulfments (Fig. 5B, C).

**Figure 5:**
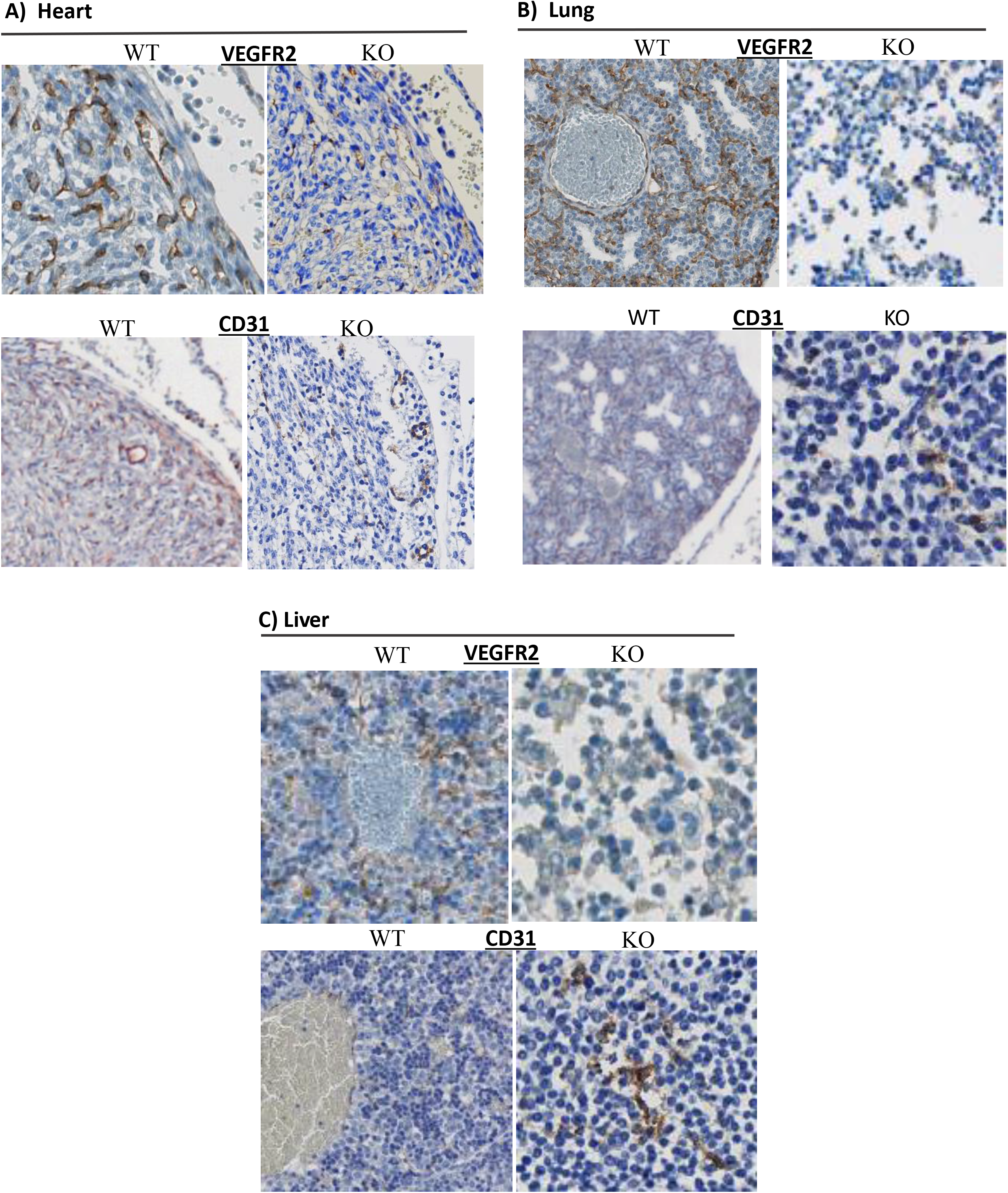
ED14 WT compared to ED14 KO: Capillary Markers. Sagittal slices approximately sequential to those stained with H&E were immunostained for CD31 or VEGFR2, which are commonly used to identify endothelial cells (ECs), but can also stain epithelial cells. A) Heart: In the WT, both markers stain ECs lining capillaries and larger blood vessels and epithelial cells. CD31 also faintly stained cardiac muscle cells. In the KO, CD31 stains what appear to be failed blood vessels near the heart’s surface, while VEGFR2 staining is less structured; scattered cells and aligned cells stain, and apparent degraded cells stain faintly in patterns suggesting failed blood vessels. B) Lung: In the WT, both markers stain ECs lining capillaries and larger blood vessels in similar patterns; VEGFR2 capillary staining is stronger, and VEGFR2-stained cells completely encircle large blood vessels. In the KO lung, CD31 moderately, and VEGFR2 faintly, stain apparent cell fragments and leaked cytoplasm, but structural patterns are no longer evident. C) Liver: In the WT, VEGFR2 stains capillaries, but few stained cells line a large blood vessel; CD31 capillary staining is less pronounced, but broken threads of stained cells line a large blood vessel. In the KO, VEGFR2 is quite faint, but scattered cells and apparent degraded cells can be seen. Oddly, CD31 signal in the KO liver was stronger than in the WT; a few scattered lines and clusters of degraded cells and possible leaked cytoplasm may be remnants of failed blood vessels.

We explored connections between blood vessels (Fig. 5) and SAMD1 (Fig. 6). At ED14 in the WT heart, anti-SAMD1 monoclonal fab-fragment (fab1961) and CD31 stained essentially identically in capillaries and epicardium, but less intensely than VEGFR2. CD31, VEGFR2, and fab1961 stained diffusely in similar locations in the lung and liver, and but fab1961 staining was weaker. In comparison, polyclonal anti-SAMD1 antibodies (GP-abs, see Methods) stained many cell types and tissues throughout the embryo, rarely resembling CD31 or VEGFR2, instead producing a range from strong cell-associated staining, to faint diffuse cytoplasmic staining, and apparent extracellular staining. fab1961 and GP-abs stained in similar patterns in the diaphragm, lung, and liver, but strikingly differently in the kidney and heart. GP-abs also stained scattered cells in the blood, root ganglia at the spine, and cells and extracellular matrix (ECM) in the brain.

**Figure 6:**
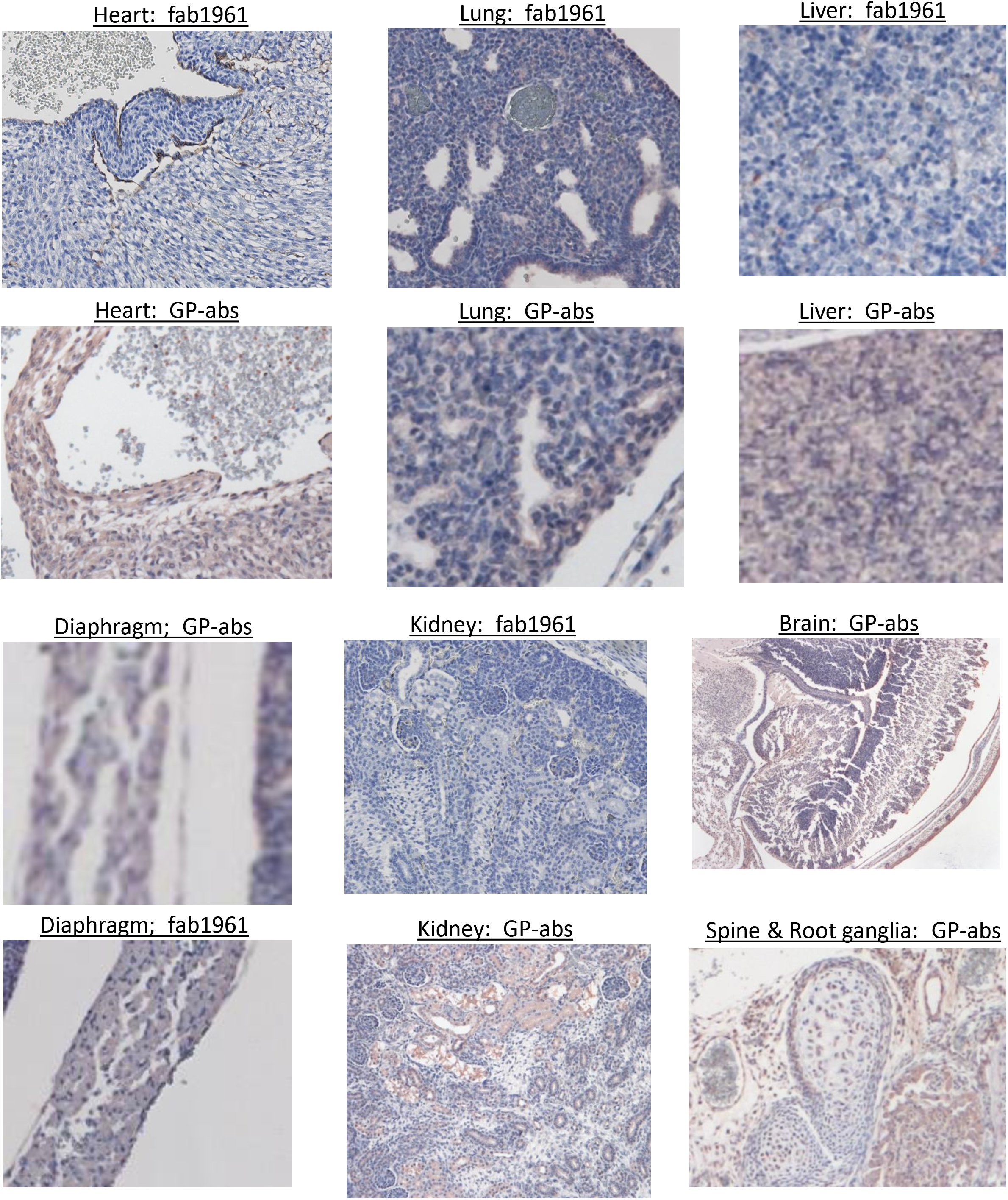
Staining differences between polyclonal and monoclonal SAMD1 probes in ED14 WT suggests variations in epitope exposure during organ development. Slices approximately sequential to those stained with H&E were immunostained for SAMD1 using polyclonal antibodies made in guinea pigs against the 3’ (C-terminus) half of rabbit SAMD1 (GP-abs) or a monoclonal fab fragment inhibitor of SAMD1/LDL binding made against full length human SAMD1 (fab1961). In the heart, fab1961 stained ECs and endocardium indistinguishably from CD31; in contrast, GP-abs stained ECs, epicardium, endocardium, myocardium nuclei and cytoplasm, and some cells in the blood. GP-abs and fab1961 show similar diffuse cytoplasmic stain in the diaphragm muscle cells. In the lung, GP-abs and fab1961 staining patterns were similar. fab1961 EC staining is fainter and more diffuse than CD31, but fab1961 epithelial staining is stronger, and also stains alveoli (Fig. 5). GP-abs and fab1961 stain the liver in similar patterns, but GP-abs stain more strongly compared to the faint fab1961 staining. fab1961 EC staining throughout the kidney is weaker than GP-abs; here fab1961 stains some only ECs, glomeruli, and tubules. GP-abs shows uniform cytoplasmic, microvilli, and occasional nuclear staining in kidney tubules. In the spine, GP-abs stain ECs, endochondral cells, root ganglia, and unidentified cells. GP-abs staining is seen throughout the brain, skull, and dermis (dermis is also seen in Fig. 7), in nuclei, cytoplasm, and apparently extracellularly. At first glance, GP-abs appear to stain promiscuously; closer examination shows numerous cells with similar morphologies that do and don’t stain, the exception being muscle cells.

Comparison of CD31 to SAMD1 staining in the ED14 KO was used to investigate possible genetic compensation for lack of SAMD1 in blood vessel development. ED14 WT dermal layer staining for CD31 or GP-abs, when compared with the ED14 KO, showed interesting similarities and differences (Fig. 7A). CD31 in the WT stained only obvious capillaries, and some rounded cells. In contrast, GP-abs stained keratinocytes, myocytes, hair follicle cells, and stained tentatively identified fibroblasts, lymphatic capillaries, immune cells, and nerve cells. Similar to CD31, SAMD1 staining in the KO marked dotted EC strands and partial circles, suggesting that both probes are evidencing failed blood vessels; SAMD1 also stained degraded hair follicles and numerous rounded cells, but cytoplasmic muscle staining was attenuated. CD31 around endochondral bone in a forepaw displayed broken strands from likely apoptotic ECs; GP-abs staining was minimal except for one short stretch on the skin (Fig. 7B). CD31 stained broken strands adjacent to ribs, and broken circles in muscle between the ribs. GP-abs staining patterns were similar but consisted of more numerous small clusters of nuclei; as above, cytoplasmic muscle staining was attenuated compared to WT (Fig. 7C).

**Figure 7:**
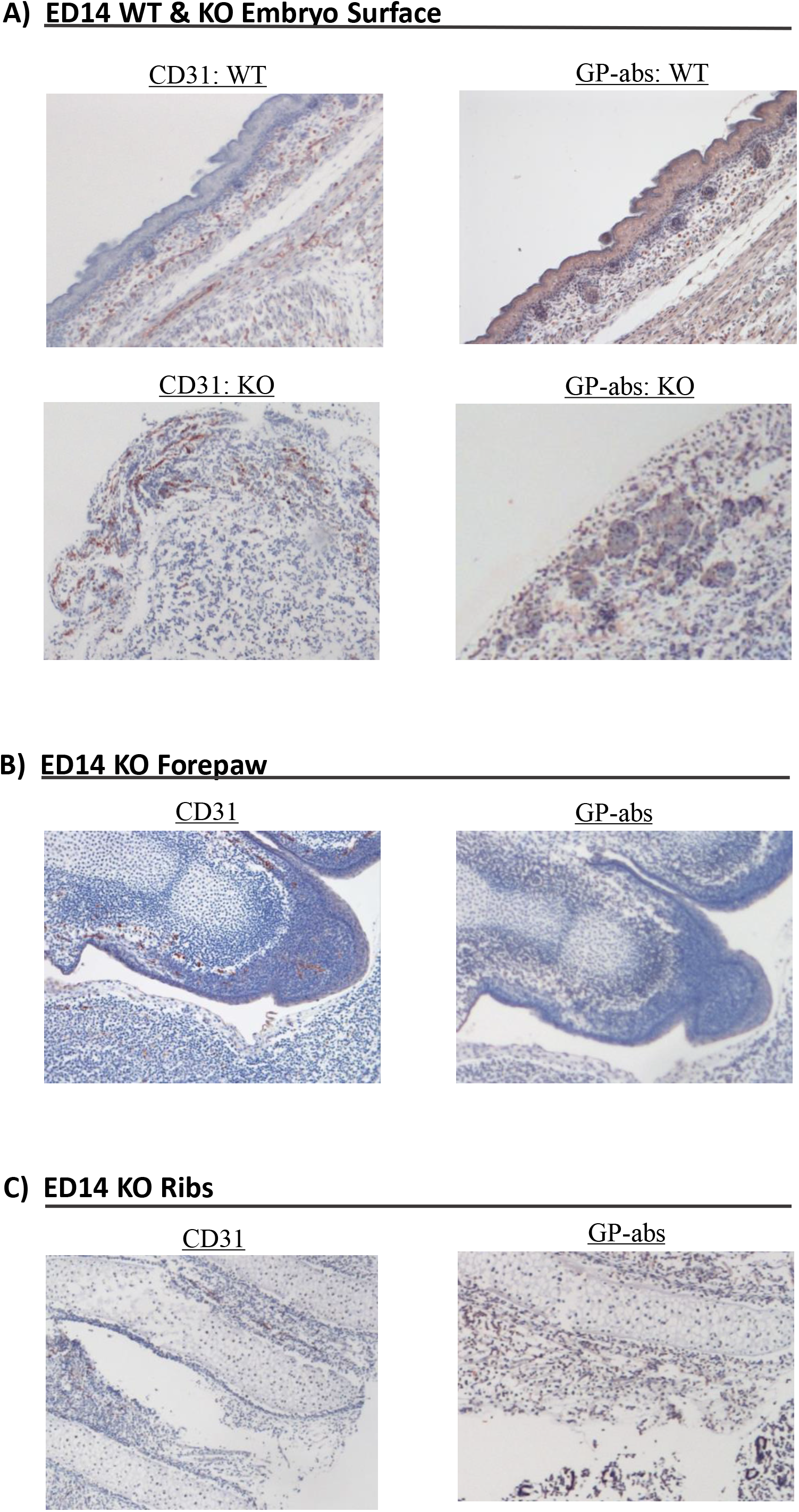
Similarities in SAMD1 and CD31 staining patterns in WT and KO mouse skin and bone suggest failed attempts to form blood vessels. A) Slices from similar areas at the embryo surface of ED14 WT and KO mice were IHC stained using GP-abs for SAMD1, or CD31 for ECs and epithelial cells. The WT has intact skin, here CD31 stains only ECs around capillaries and small vessels, and some rounded cells. In the WT, uniform SAMD1 stain appears in keratinocytes, extracellular material in hair follicles, and muscle cell cytoplasm. Typical elongated ECs around capillaries are not apparent with SAMD1, but elongated cells run through what may be an adipose layer. Two cells in this layer have a dendrite morphology; one stains for SAMD1, the other does not. Some rounded cells in the dermis, many of which have eccentric nuclei stain, other rounded cells do not stain. The KO epidermis is missing, and the embryo surface is disorganized, but in patterns reminiscent of the WT. CD31 in the KO stains cells in patterns that suggest failed capillaries, and GP-abs stain scattered nuclei in the dermis, some of which form dotted strands or broken circles, extracellular material in hair follicles, and numerous possible immune cells. B) In a KO forepaw that was relatively well formed, CD31 stained in the perichondrium, and in a few places had invaded the cartilage, but obvious capillaries had not formed; faint diffuse stain appears in what should become skin. GP-abs staining was almost absent, except for a strand where skin would form. C) Ribs in the KO also stained for CD31 in broken lines, but endothelial cartilage invasion was not apparent; GP-abs appeared to stain similar lines, but extensive staining of skeletal muscle cytoplasm and nuclei, although weaker than in the WT, obscures these signals.

### SAMD1−/+ Phenotype Compared to Wild Type

SAMD1 mRNA levels were about 21% higher in MEFs from WT compared to MEFs from HETs (Fig. 1). Genotyping 3 weeks after birth showed the number of observed HETs was significantly lower than the expected number, both when SAMD1+/− mice were crossed with SAMD1+/− mice (p<0.00274; 1:2 ratio), and when SAMD1+/+ mice were crossed with SAMD1+/− mice (p<0.00323; 1:1 ratio) (Fig. 1). However, the majority of HETs survived until sacrifice at 52 weeks. Multiple characteristics were measured to test for differences between adult WTs and adult HETs (N=5). Corticosterone, aldosterone, and angiotensin II levels were meaningfully increased in the HET mice, and testosterone levels trended higher (Supplementary Fig 2). Longitudinal body weight analysis on regular chow revealed that male and female WT mice weighed more than HET mice. On high fat diet (HFD), fat depots from WT weighed twice as much as fat depots from HETs, after correction for body weight. The HETs had increased glucose disposal during the oral glucose tolerance test, lower baseline fasted glucose levels, and unchanged insulin levels following the HFD challenge. The HETs’ increased glucose disposal and hypercortisolemia are consistent with lower adiposity and muscle weight. There was a relative increase in the percentage of cholesterol in VLDL particles, and a trend towards an increased percentage of LDL with a corresponding decrease in the percentage of cholesterol in HDL particles after the high fat diet challenge (HFD) in the HET mice, but no effect on the total cholesterol level was observed. The HETs exhibited meaningfully lower serum alkaline phosphatase and a trend (p=0.092) towards lower serum thyroxine (T4) levels. Higher inorganic phosphorus levels were found in the serum. The prothrombin time was meaningfully lower in the HET as compared to WT mice, while thrombin-antithrombin levels tended to be higher in the HET as compared to WT mice (p=0.069). The urine creatinine levels tended to be higher in the HET as compared to WT mice (p=0.075).

The HET mice appeared to have an increased response in the OVA-induced pulmonary inflammation model as evidenced from a trend towards higher number of Gr1+ cells in the broncho-alveolar lavage fluids (p=0.0798). In addition, the total serum IgE levels (p=0.122) and levels of IL-10 in the lung (p=0.1288) appeared to be increased in the HET mice in this model. However, the HET mice produced meaningfully less IgG1 after secondary stimulation with OVA in the *in vivo* antibody production assay and the numbers of eosinophils, monocytes and basophils (P<0.10) were decreased in the peripheral blood of the HET mice. *In vitro* LPS challenge of monocytes revealed meaningfully lower IFN-γ, IL-10, TNF-α, and mKC levels in the HET as compared to WT.

There were no meaningful differences in B-cell proliferation, bleeding time, collagen induced platelet aggregation, cytokine production by T-cells, sexual behavior (male or female), formalin induced pain, von Frey mechanical pain, tail suspension test, pre-pulse index, rotarod, monocyte infiltration, free fatty acids, liver or muscle glycogen, or behavior (body temperature, heart rate, pupil reflex, respiration, body position, limb grasping, locomotor activity, tail pinch, toe pinch, body tone, gait, grip strength). Serum and urine electrolytes did not differ between the HET and WT mice.

CD31 and SAMD1 staining in an ED14 HET were not different from the WT (Supplementary Fig. 3). Atherosclerotic lesion size and appearance were similar when comparing SAMD1+/− apoE−/− mice to the apoE−/− mice control group after 35 weeks on a modified western diet (n=12).

## Discussion

### SAMD1−/− Mice

In this study, we investigated the effects of ablation of the SAMD1 gene in mice. We show that SAMD1 is needed for fetal angiogenesis, and is likely involved in other aspects of embryonic development.

In mice, a simple circulatory loop is developed by ED8; heartbeats begin around ED8.5, and circulatory flow of primitive erythroblasts from ED8.5-ED9.5 is necessary to remodel early vessels into a network of branched endothelial tubes [20]. Atrial function is necessary for embryonic development beyond ED10.5 [21]. Mutations which affect ECs tend to be embryonic lethal by ED10.5 [20]. After initial vasculogenesis, hypoxia induced signaling recruits mesenchymal progenitors that contact EC tubes and between ED10.5 and ED13.5; the timings and sources of the VSMC progenitor cells varies by vessel, tissue, and organ; in the heart, VSMC markers emerge in cells adjacent to ECs around ED14.5 [22]. SMCs reorient to wrap circumferentially around the developing vessel; a second layer is added around ED 13.5 [23]. These changes also occur in yolk sac vasculature [24]; around ED14, normally developing blood vessels emerge in the yolk sac and embryo [20]. Lack of, or defective, SMCs cause vascular regression and pruning by ED12, and tend to cause death between ED12 and ED17 [25].

Hypoxia is necessary for cell proliferation and embryonic morphogenesis up until a point at which hypoxia impedes development and causes apoptosis [26]. Hypoxic apoptosis is delayed in hypertrophic chondrocytes [27], radial glia, and neural precursor cells [28].

Absence of SAMD1 seemed to have little effect on development prior to ED12, but at ED14, differences were obvious to the naked eye. Most KO tissues and organs were recognizable with IHC microscopy; for example, cardiac and skeletal muscle cells were obvious, but incomplete development and/or necrotic degradation made some identifications difficult (Supplementary Fig. 4). Scattered lines and small accumulations of nucleated and enucleated RBCs demonstrate that vasculogenesis, early hematopoiesis, and functional heartbeat had occurred. Staining patterns of the endothelial markers CD31 and VEGFR2 in the WT compared to the KO provide further evidence of early endothelial tube formation, but suggest that endothelial tubes degraded before developing into capillary networks, as inferred from: a) the lack of capillaries and larger vessels in the lungs; b) the fact that the few RBCs seen were not contained in vessels; and c) the stained strands and broken circles of probable apoptotic ECs in the heart; at the embryo surface; and adjacent to developing bones. These data suggest that angiogenesis failed subsequent to vasculogenesis. Lack of angiogenesis would prevent relief from hypoxia. Extended hypoxia is the likely cause of: the apoptosis/necrosis apparent in many locations; the hypertrophic chondrocytes in the ribs; and the enlarged brain in the ED15.5 mouse. Fluid leakage due to lack of blood vessels likely produced the edema.

SAMD1’s role in successful embryonic vessel development may involve ECs and VSMC progenitors. Monoclonal anti-SAMD1 fab1961 and CD31 essentially identically stained both ECs, and endocardial cells (which are a source of VSMC progenitor cells [22]), in the ED14 WT heart. In earlier work, we showed that SAMD1 stains VSMCs and ECs in adult mice, such that changes in staining patterns appear to correlate with these cells’ differentiation and dedifferentiation, and that SAMD1 may be involved in VSMC and EC processing/binding of cholesterol, possibly for use in blood vessel repair [3]. In cell culture, SAMD1 was shown to be involved in epigenetic control of VSMC migration, proliferation, and/or differentiation [5], but VSMCs, at least in the heart, are thought to develop after ED14 [22].

The presence of SAMD1 mRNA throughout the WT embryo and yolk sack, at levels that varied during gestation, suggests that SAMD1 is used beyond vessel development. As a possible example, the single highest spike in SAMD1 mRNA was seen in the head at ED12; lack of this may have prevented differentiation of the cephalic mesoderm cells that form the skull vault [29], causing the exencephaly seen at ED15.5 (Fig. 3). This is supported by a recent finding that ablation of SAMD1 in embryonic stem cells resulted in impaired differentiation [6].

Epigenetic proteins are found in cytoplasm and nuclei, and are widely expressed during embryonic development [30]. Staining for SAMD1 with polyclonal GP-abs roughly correlates with reports of polycomb staining patterns seen in the liver [30], kidney [31], and brain [32]. This suggests the GP-abs IHC data is likely to be largely valid, despite the risk of crossreactivity. Further, polyclonal and monoclonal SAMD1 staining patterns are very similar in some tissues (liver, diaphragm) and quite different in others (heart, kidney) (Fig. 6), (Supplementary Fig. 5). This observation supports the validity of the polyclonal staining, while suggesting that the monoclonal-targeted epitope may not be consistently available.

Interestingly, SAMD1, and the EC markers CD31 and VEGFR2, all stained what appear to be EC fragments and apoptotic/necrotic cells in the ED14 KO, in locations and patterns that suggest failed blood vessels (Fig 7), (Supplementary Fig. 5). SAMD1 staining in the KO is not unexpected, since production of compensating proteins, which typically have domain similarities with a knocked-out protein, allow at least partial rescue of a developing embryo [33]. We speculate that anti-SAMD1 antibodies are binding to compensation proteins generated by the KO’s attempts to induce VSMC proliferation and initiate capillary EC investment with VSMCs. Exploration of differential gene expression between WT and KO could provide useful data, as might conditional SAMD1 knockouts, and using anti-SAMD1 abs as bait for potential compensating proteins expressed by the KO.

### SAMD1 Heterozygous Mice

Mutations that result in heterozygous absence of a gene often do not have uniform effects across a population; differences in penetrance, i.e. whether or not they exhibit signs of the mutation, and variable expression i.e. to what extent they exhibit one or more signs of the mutation, are common; the mechanisms can be environmental, epigenetic, synergistic, and multigenic [34]. Variable expression and penetrance are likely explanations for the approximately 30% postnatal HET mortality by 3 weeks of age, but causes of death were not available for these mice. Surviving HETs had 18% lower SAMD1 levels, and some mice had substantially increased aldosterone, corticosterone, angiotensin II, and testosterone levels (Supplementary Fig. 2). Steroidogenesis and transformation of cholesterol to oxysterols occurs in many cell types [35], so it is possible either that SAMD1 directly regulates hormone production, or that lower SAMD1 levels cause multigenic compensatory changes in steroid expression. The HET’s reduced adiposity and muscle mass may be related to the steroidal changes. Insufficient data was available to hypothesize about connections between reduced SAMD1 and the other various trends noted in the surviving HETs.

Heterozygous knockout of SAMD1 did not have a significant effect on lesion size when comparing apoE−/− control mice and SAMD1+/− apoE−/− mice (N=12) after 35 weeks on modified western diet. It appears that sufficient SAMD1 was expressed by the surviving mice to allow essentially normal function; thus, no connections to atherosclerosis can be inferred. As is typical in mouse atherosclerosis models, lesion areas varied widely within each group. The study was powered to account for this, but it is possible that a much larger study could find a dose response.

## Conclusions

This is the first report on ablation of SAMD1 in mice; embryonic lethality was observed, with failure of fetal angiogenesis a prominent result, possibly through non-development of VSMCs. Proliferation, migration, and/or differentiation at least of cephalic mesoderm cells were also likely affected. Embryonic lethality may be primarily caused by the myriad effects of lack of blood vessels, but, given SAMD1’s proposed epigenetic functions, numerous other factors may be involved. 3-week postnatal HET mortality rates imply that some individuals cannot survive the effects of reduced SAMD1. This study provided no data on SAMD1’s possible role in atherosclerosis; it would be interesting to create conditional SAMD1 knockouts to compare atherosclerotic lesion size in apoE−/− mouse or LDLR−/− mouse models of atherosclerosis after at least 35 weeks on western diet.

## Materials and Methods

### SAMD1 Gene Inactivation

SAMD1 knockout mice were genetically engineered by Pfizer (Groton, CT). To generate the SAMD1 targeting vector, recombineering [36] was used to replace 2396 bp of the mouse SAMD1 (Accession ID D3YXK1) gene encompassing the 3’ 615 bp of exon 1, exons 2-4 and all of exon 5 except for the 3’ 83 bps with a neomycin phosphotransferase cassette in a 9433 bp genomic subclone obtained from a C57BL/6J BAC (RP23-128H6; Invitrogen). The vector was introduced into Bruce4 (mixed C57Bl/6N:C57BL/6J) mouse embryonic stem cells [37] using standard homologous recombination techniques [38]. Southern analysis was used to identify correctly targeted mouse embryonic stem cell clones. Chimeric mice were generated by injection of the targeted ES cells into Balb/c blastocysts [39]. Chimeric mice were bred with C57Bl/6J mice to produce F1 heterozygotes. Germline transmission was confirmed by PCR analysis (SAMD1g1: 5’-CCAAACCCCTCTTCAGTTCA-3’; SAMD1g2: 5’-GCCGTAGCTATTCTGCCTCA-3’; dNEO2: 5’-ACATAGCGTTGGCTACCCGTGATA-3’). F1 heterozygous males and females were mated to produce F2 mice. The colony was maintained by intercrossing under specific pathogen free conditions with unrestricted access to food and water. A separate colony was similarly bred from the above mice by the Brigham & Women’s Hospital (Boston, MA)

### RNA and cDNA Measurements

RNA and cDNA measurements were performed at the Brigham & Women’s hospital (Boston, MA). Synthesis of cDNA (ThermoScript RT-PCR System (Invitrogen Cat 11146-016)). RT-PCR was performed in a MyiQ single-color real-time PCR system (Bio-Rad Laboratories, Inc., Hercules, CA). Total ribonucleic acid (RNA) from 250,000 cells was reverse transcribed by Superscript II (Invitrogen) according to the manufacturer’s instructions. Quantitative PCR was performed with SYBR green PCR mix (primers for SAMD1 from QIAGEN N.V.) and analysis performed with StepOne Software (ThermoFisher, Applied Biosystems). Levels of mRNA were normalized to glyceraldehyde 3-phosphate dehydrogenase (GAPDH) mRNA levels. SAMD1 sample and controls cDNAs were synthesized (ABI high-capacity cDNA Archive Catalog No. 4387406). Total 2ugs RNA/100ul cDNA synthesis reaction and 1ul cDNA per Taqman Reaction. Mouse SamD1 Assay – Mm01261650_g1. Mouse GAPDH endogenous control-(Applied Biosystem 4352339E. Catalog No. 43-523-39E). MEF mRNA studies were performed at the Brigham & Women’s hospital (Boston, MA). MEFs were isolated as described [40]. Embryos from WT, SAMD1+/−, and SAMD1−/− mice were isolated between ED12 and ED18. Embryos were minced and trypsinized after removal of most internal organs, heads, tails, and limbs were removed, then seeded into T-75 cell culture dishes in 15 mL of complete MEF media.

### Phenotype Studies

All behavioral, physical, hormonal, and response phenotype comparisons of WTs to HETs were performed by Caliper Life Sciences (Xenogen Biosciences, Cranbury, NJ).

### Immunohistochemistry

Fully human monoclonal fab-fragments (fabs) to full-length human SAMD1 were made by Biosite (Biosite/Alere/Abbott, San Diego, CA) using phage display; Biosite selected fab1961 by ELISA for affinity to full-length human SAMD1, and ability to inhibit SAMD1/LDL binding. Full-length human SAMD1 was produced by Cell & Molecular Technologies (CMT, Phillipsburg, NJ, now Invitrogen, Carlsbad, CA) in HEK293 cells: the human SAMD1 coding sequence (1617bp) was redesigned such that the codon usage was optimized without changing the amino acid sequence. Polyclonal antibodies were generated in guinea pigs against a rabbit SAMD1 fragment expressed in e coli that started at a.a. 238 and continuing to the 3’ end (GP-abs), as previously described [16]. Antibodies to CD31 and VEGFR2 were purchased (Abcam, Cambridge, UK). Immunohistochemical staining procedures were performed by Pfizer (Groton, CT) and at the Brigham & Women’s hospital (Boston, MA).

### Lesion Measurement

Lesion studies were performed at the Brigham & Women’s hospital (Boston, MA). SAMD1+/− apoE−/− and apoE−/− mice (N=12) were fed a HCHF diet for 35 weeks to induce spontaneous aortic lesions. Lesion area was calculated by using Image J software to determine the red ORO *enface* stained areas of the main trunks of the artery from the aortic root to the iliac bifurcation, as previously described [41].

## Acknowledgements

We thank Dr. Andrew Lichtman (Brigham & Women’s Hospital, Boston, MA) for guiding discussions, critical inputs, unwavering intellectual support, and for managing members of his lab throughout this research. We thank Dr. Margaret Tarrio (Brigham & Women’s Hospital, Boston, MA), for breeding, raising, performing microscopy, and IHC studies on KO mice; for performing various assays; and for mouse treatment studies. We thank Dr. Gunars Valkirs (Biosite /Alere/Abbott, San Diego, CA) for guiding the development and testing of fab fragment inhibitors of SAMD1/LDL binding.

---

**Supplementary Figure 1:**
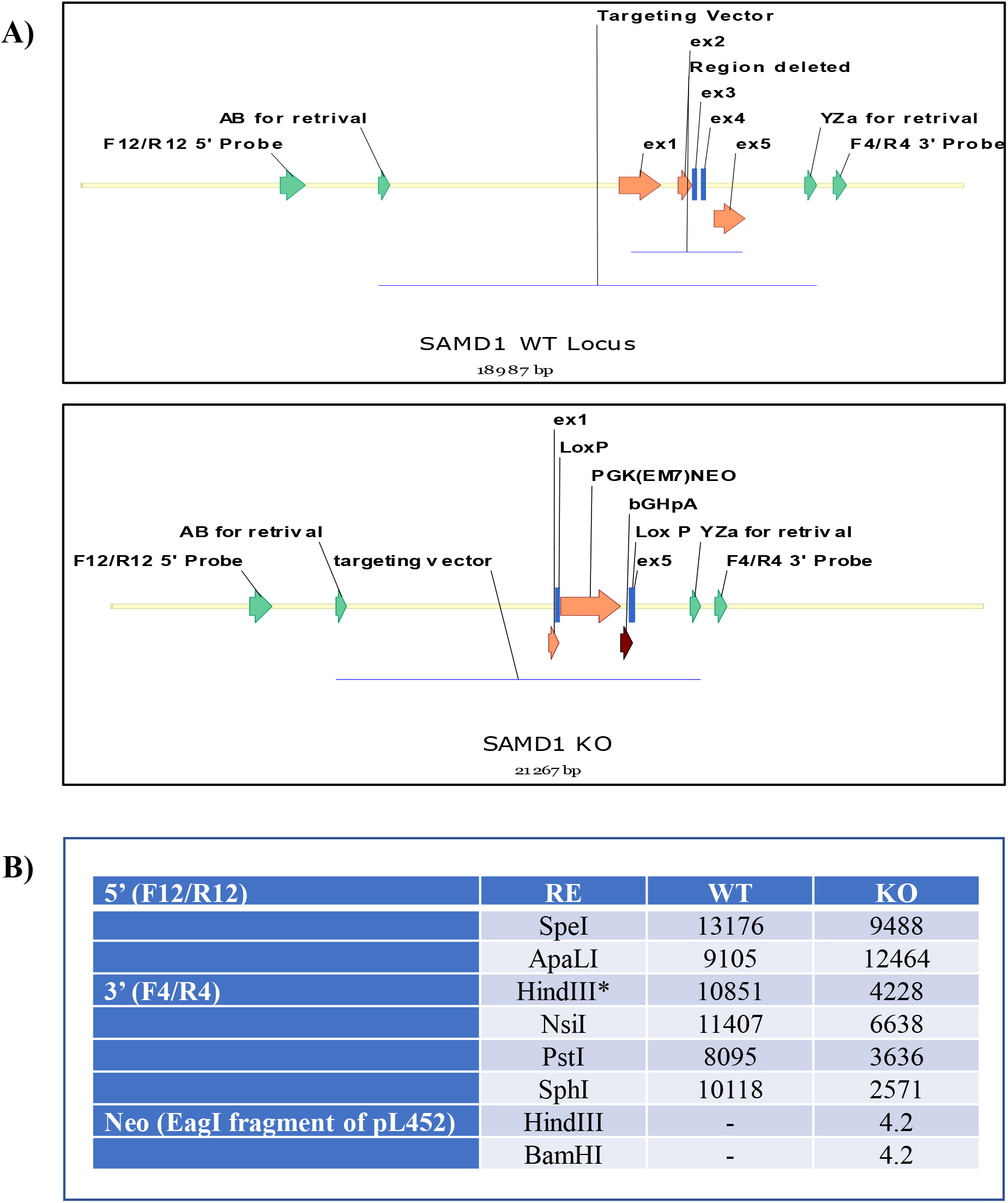
Knockout targeting and validation. The first 256 bp of the gene were left intact, however, only 63 bp of the 5’ exon were left intact; since this is past the stop codon for the protein there are no additional 3’ amino acids. A) The targeting vector for SAMD1 gene knockout replaces 2396 bp of the SAMD1 gene encompassing the 3’ 615 bp of exon 1, exons 2-4 and all of exon 5 except for the 3’ 83 bps with a neomycin phosphotransferase cassette in a 9433 bp genomic subclone. The primers to generate the Southern probes that validated targeting are: SAMD1F4,5’ -AACCAGGTTCCATCTGCATC-3’: SAMD1F4,5’-GAACCACCTGCATCTGTCCT-3’ SAMD112, 5’-CACTTGGGGATGTGGTCATATTCA-3’: SAMD112, 5’-CTCGTAGATGAGGACTCCGAGAGC-3’ B) Validation strategy used restriction sites SpeI, ApaLI, HindIII, NsiI, PstI, SphI, and BamHI

**Supplementary Figure 2.**
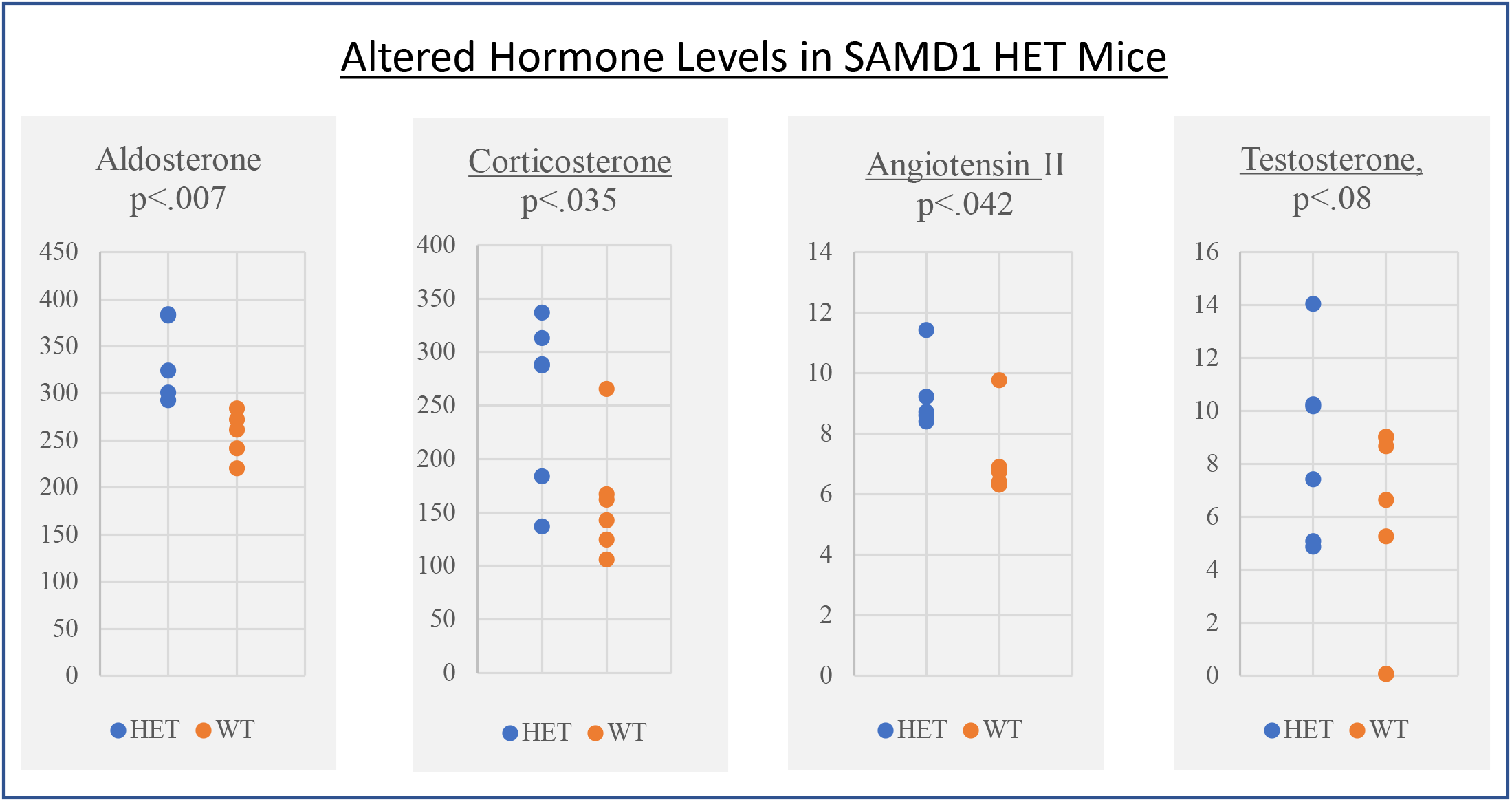
Increased hormone levels may be related to failure to thrive. Increased levels of the hormones aldosterone, corticosterone, angiotensin II, and testosterone were found in SAMD1 HET mice. The few HETs with the highest hormone levels may be related to the failure to thrive. Although the p-value for increased testosterone does not reach statistical significance due to the size of the scatter, the data strongly suggests a trend.

**Supplementary Figure 3.**
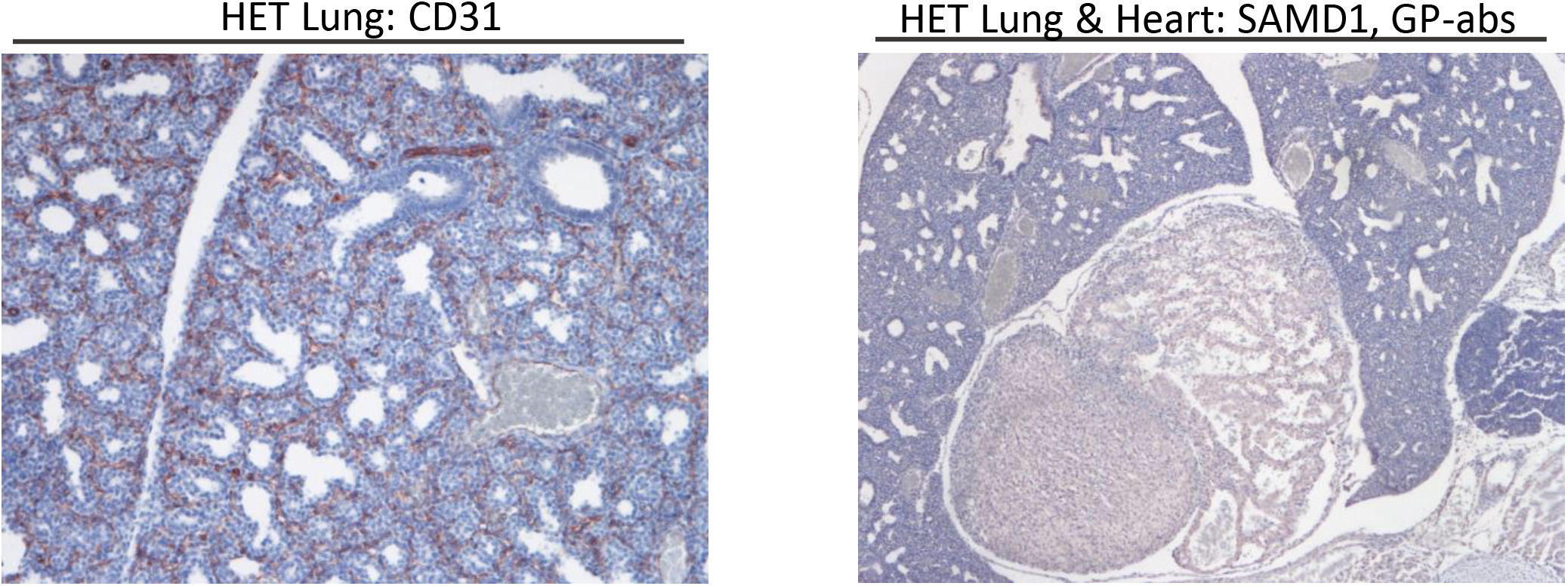
CD31 and SAMD1 staining of a HET lung do not appear materially different from the WT lung stained for CD31 or SAMD1. Two IHC images from an ED14 HET mouse are included to show that organ formation in at least one ED14 HET appeared normal.

**Supplementary Figure 4.**
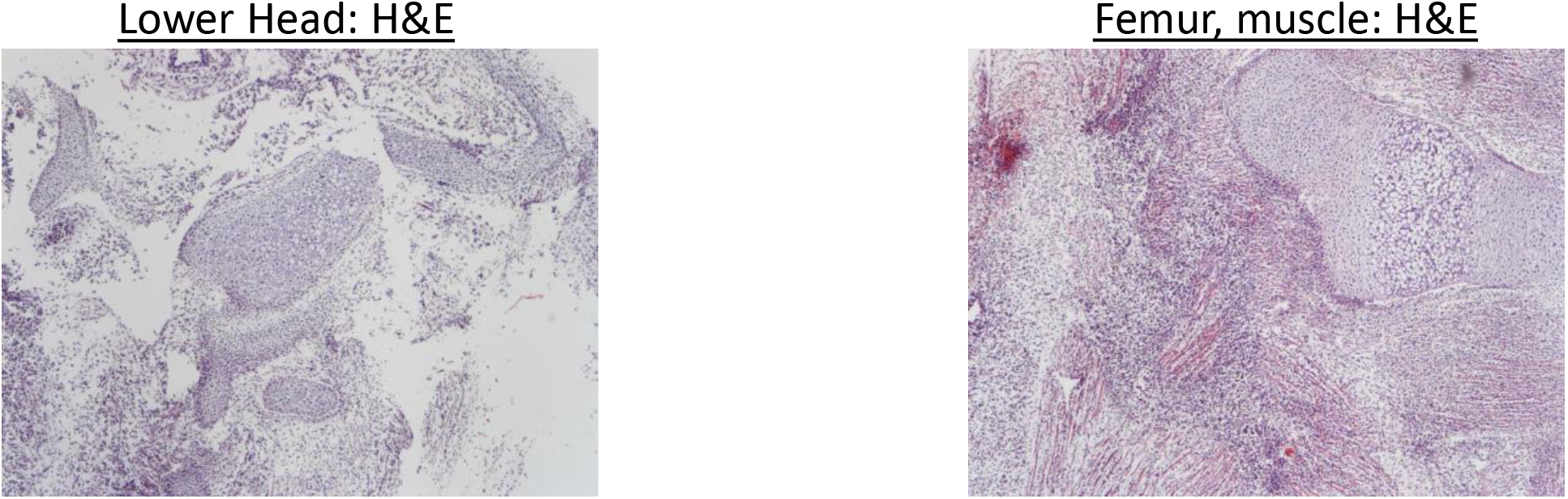
Some tissues are underdeveloped or extensively degraded.

**Supplementary Figure 5.**
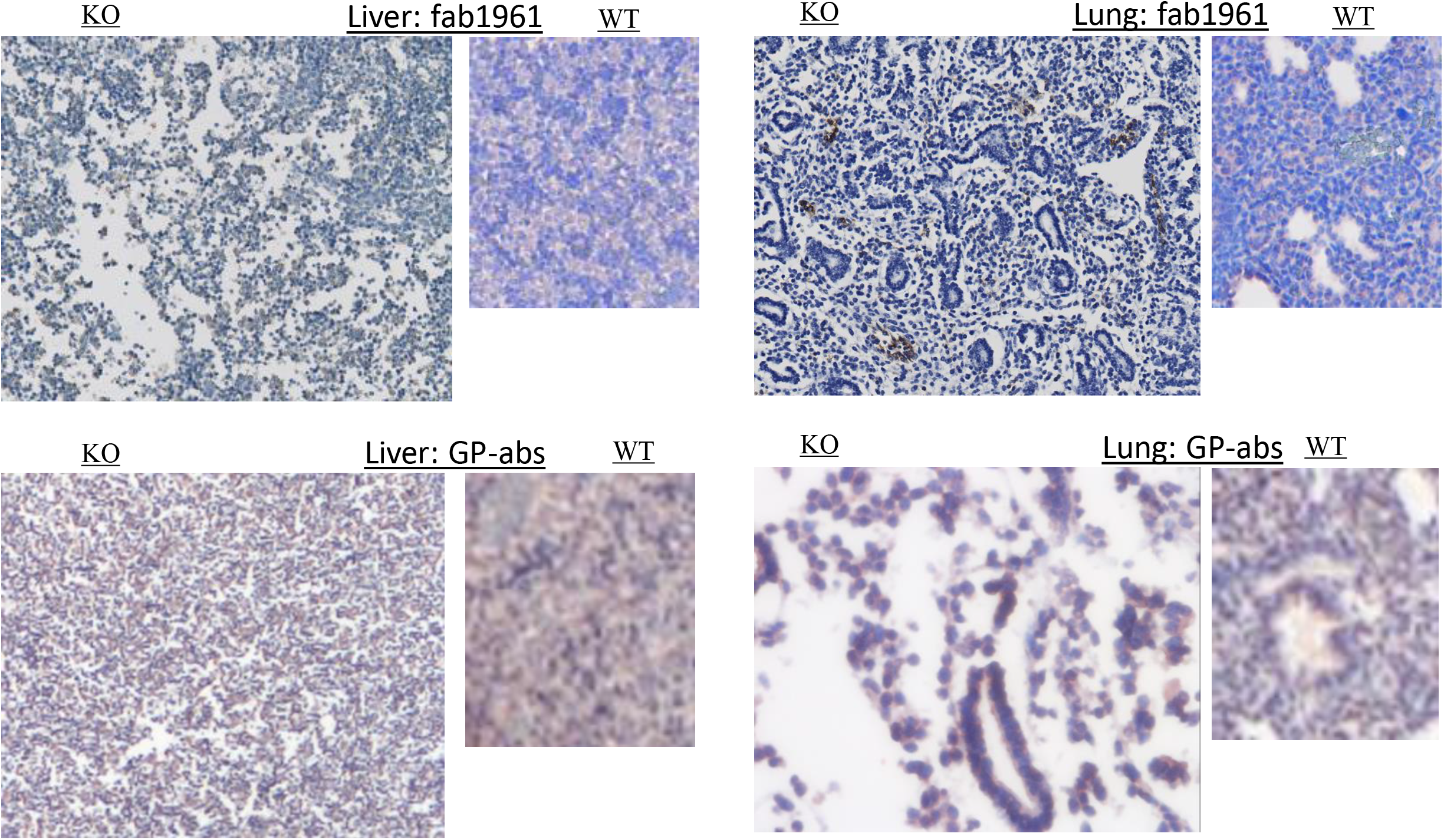
ED14 KO embryo slices were stained with polyclonal GP-abs or monoclonal fab1961 in an attempt to show presence of possible compensating proteins. Monoclonal and polyclonal SAMD1 abs in ED14 mice stain similarly in some tissues, differently in other tissues; patterns between KO and WT for each stain are roughly similar. In the KO liver, GP-abs stain uniformly in locations typical of capillaries; some staining appears cell-associated, some appears extracellular, and some may be cell fragments; fab1961 stains similarly, but much more faintly. In the KO lung, alveoli stained with GP-abs, but fab1961 stained possible blood vessels and some scattered clusters of cells that were not clearly associated with recognizable structures.

